# Gut microbiota of *Drosophila subobscura* contributes to its heat tolerance but is sensitive to transient thermal stress

**DOI:** 10.1101/2021.01.08.425860

**Authors:** Angélica Jaramillo, Luis E. Castañeda

**Affiliations:** Programa de Genética Humana, Instituto de Ciencias Biomédicas, Facultad de Medicina, Universidad de Chile, Santiago, Chile

**Author notes:** Corresponding author: Luis E. Castañeda.

**Keywords:** bacterial microbiota, climate change, heat stress, stress resilience, thermal resistance

## Abstract

Global warming impacts the animal’s fitness, leading to increasing extinction risk in ectotherm species. Gut microbiota can contribute to host physiology leading to an increase of resistance to abiotic stress conditions. Temperature has profound effects on ectotherms and gut microbiota can influence cold and heat tolerance of ectotherm species. Additionally, the gut microbiota is sensitive to environmental temperature, which induces changes in its composition and diversity. Here, we investigated the role of the gut microbiota on heat tolerance of *Drosophila subobscura*, comparing the knockdown time between conventional and axenic, and we also assessed the impact of heat stress on the diversity and community structure of the gut microbiota, comparing non-stressed and heat-stressed flies. Our findings provide evidence that gut microbiota influences heat tolerance, and that heat stress modifies the gut microbiota at taxonomical and structural levels. These results demonstrate that gut microbiota contributes to heat tolerance but it is also highly sensitive to transient heat stress, which could have important effects on host fitness, population risk extinction, and vulnerability of ectotherms to current and future climatic conditions.

## Introduction

Gut microbiota influences multiple features of the host’s biology, including nutrient acquisition, immune response, metabolism, behavior, and life-history traits (Broderick and Lemaitre, 2012; Douglas, 2018a; Hoye and Fenton, 2018). In general, gut microbiota influences phenotypic variation exhibited by host organisms, which can contribute speeding up the adaptive response under changing and fluctuating environments (Alberdi et al., 2016; Macke et al., 2017; Romano, 2017). However, environmental variability, ranging from benign to stressful conditions, could impact the composition and diversity of the gut microbiota, altering its contribution to host phenotypic variability and modifying the functional relationship between hosts and the gut microbiota (Sepulveda and Moeller, 2020).

Among multiple environmental factors, temperature has profound effects on physiology, behavior, and performance of ectotherms, because body temperature of ectotherms is influenced by the environmental temperature (Angilletta, 2009). The ongoing climate change is expected to impose strong selection pressures on natural populations of ectotherms (Hoffmann and Sgrò, 2011), and ectotherms can display genetic and non-genetic mechanisms that contribute to host thermal tolerance (Kokou et al., 2018). Indeed, recent evidence demonstrates that microbiota impacts on thermal performance of ectotherm species (Renoz et al., 2019; Sepulveda and Moeller, 2020). For instance, obligatory endosymbionts contribute to aphid performance at high temperatures (Dunbar et al., 2007; Zhang et al., 2019), whereas facultative endosymbionts also confer tolerance to high temperature in aphids (Montllor et al., 2002; Russell and Moran, 2006) and *Drosophila* (Gruntenko et al., 2017). Additionally, it has been demonstrated that microbiota also influences cold and heat tolerance of ectotherm species (Ziegler et al., 2017; Henry and Colinet, 2018; Kokou et al., 2018; Moghadam et al., 2018; Raza et al., 2020).

On the other hand, several studies have also explored the impact of temperature on host microbiota, indicating that gut microbiota is sensitive to environmental temperature (Wernegreen, 2012; Sepulveda and Moeller, 2020). Temperature induces changes in the composition and diversity of the gut microbiota, which could have important consequences on host phenotype and fitness (Wernegreen, 2012; Alberdi et al., 2016). For example, small ectotherms reared at high temperatures show an increase in the abundance of bacteria belonging to the Proteobacteria phylum (Li et al., 2018; Moghadam et al., 2018; Horváthová et al., 2019). Indeed, *Drosophila melanogaster* flies acclimated in warm conditions showed a higher abundance of *Acetobacter* bacteria (Proteobacteria) and a lower abundance of *Leuconostoc* bacteria (Firmicutes) in comparison to cold-acclimated flies (Moghadam *et al*., 2018). On the other hand, several studies have demonstrated that bacterial diversity and richness decrease when hosts are exposed to warm conditions (Kokou et al., 2018; Moghadam et al., 2018).

In order to investigate the role of the gut microbiota on heat tolerance of *D. subobscura*, in the present study, we compared the heat tolerance between conventional (non-manipulated) and axenic (germ-free) flies across different heat stressful temperatures. Additionally, we assessed the impact of heat stress on the diversity and community structure of the gut microbiota of *D. subobscura*, comparing non-stressed and heat-stressed flies. To study the gut microbiota of *D. subobscura*, we used a metabarcoding approach based on the amplicon sequencing of the bacterial 16S rRNA gene.

## Material and Methods

### *Drosophila* sampling and maintenance

Adult *D. subobscura* flies were collected at the locality of Valdivia (southern Chile: 39° 48’ S, 73° 14’ W) and separated by sex. Females were individually placed in plastic vials with David’s killed-yeast *Drosophila* medium (David 1962) to establish isofemale lines. At the next generation, 100 isofemale lines were randomly selected, and adult flies were dumped into an acrylic cage to set up one large outbred population, which was maintained in a climatic chamber (Bioref, Pitec, Chile) at 21 ± 1 °C and a photoperiod 12L:12D. Maintaining conditions were similar along with the experiments and the population cage was maintained on a discrete generation, controlled larval density regime (Castañeda et al., 2015).

### Preparation of axenic and conventional flies

Axenic (germ-free) flies were obtained by using dechorionated eggs (Koyle et al., 2016). Eggs (≤ 18 h-old) were collected from Petri dishes containing fly media placed within the population cage and dechorionated as follow: 3 washes with 0.5% hypochlorite sodium solution per 2 min each wash, 3 washes with 70% ethanol solution per 2 min each wash, and 3 washes with autoclaved water per 2 min each wash. Dechorionated eggs were transferred to 50-ml Falcon tubes containing autoclaved *Drosophila* media at a density of 50 eggs/tube. The procedure to obtain axenic flies was performed under sterile conditions in a flow laminar chamber. For conventional (non-manipulated microbiota) flies, eggs were collected from the same Petri dishes used previously, washed four times with autoclaved water, and transferred to 50-ml Falcon tubes containing autoclaved *Drosophila* media at a density of 50 eggs/tube.

Elimination of bacteria in axenic flies was corroborated by testing the amplification of bacterial DNA. Medium samples and 10 flies were randomly collected from tubes containing axenic flies. From both types of samples, DNA was extracted using the GeneJet kit (Thermo-Fisher) following the protocol to extract DNA from negative-Gram and positive-Gram bacteria. Then, a PCR was performed to amplify the bacterial DNA using specific primers for the 16S rRNA gene: 341F (5’-CCT ACG GGN GGC WGC AG-3’) and 805R (5’-GGA CTA CHV GGG TWT CTA AR-3’) (Fadrosh et al., 2014). The PCR mix contained 0.02 U DNA polymerase (Invitrogen), 1X PCR buffer, 0.2 mM dNTPs, 1 μM of each primer, 0.5 μM MgCl_2_, and 0.5 μl template DNA. PCR cycle conditions were set up following the recommendations of Caporaso et al. (2010b): denaturation at 94 °C for 3 min; 35 amplification cycles at 94 °C for 45 s, 52 °C for 1 min and 72 °C for 70 s; and a final extension at 72 °C for 10 min. PCR products were loaded on a 2% agarose gel stained with Sybr Safe (Invitrogen). DNA extractions from conventional flies were used as bacteria-positive controls. Thus, effective bacterial elimination was considered effective when any amplification band was visualized in the agarose gel.

### Heat tolerance of axenic and conventional flies

Axenic and conventional virgin flies of both sexes at the age of 4 days-old were individually placed in capped 5-mL glass vials, which were attached to a rack with capacity to contain 60 capped vials. In each rack, we placed 15 axenic females, 15 axenic males, 15 conventional females, and 15 conventional males. Each rack was immersed in a water tank at a specific static temperature: 35, 36, 37, and 38°C. Temperature (± 0.1 °C) was controlled by a heating unit (Model ED, Julabo Labortechnik, Seelbach, Germany). Each static assay was photographed using a high-resolution camera (D5100, Nikon, Tokyo, Japan) and photos were taken every 3 sec. Photos for each assay were collated in a video file, which was visualized to score the knockdown time (i.e. time at which each fly ceased to move (Castañeda et al., 2019).

### Heat stress exposure

Petri dishes with fly medium were placed within the population cage for collecting eggs. Eggs (≤ 18 h-old) were transferred to vials at a density of 40 eggs/vial. After eclosion, virgin flies were separated by sex and transferred to new vials. At the age of 4 days, 100 females and 100 males were transferred to empty vials at a density of 25 flies/vial, and vials were closed with moistened stoppers to avoid fly desiccation. Vials were split into two groups: non-stressed and heat-stressed flies. Non-stressed flies were transferred to a climatic chamber (Bioref, Pitec, Chile) at 21 ± 1 °C for 3 h; whereas heat-stressed flies were placed in a water bath at 34 °C for 1 h; temperature (± 0.1 °C) was controlled by a heating unit (Model ED, Julabo Labortechnik, Seelbach, Germany). This temperature was chosen because it has been previously used to induce thermal stress in *D. melanogaster* (Hoffmann et al., 2003) and *D. subobscura* (Calabria et al., 2012). Then, heat-stressed flies were transferred to a climatic chamber (Bioref, Pitec, Chile) at 21 ± 1 °C for 2 h for recovery from heat stress (no fly died after stress).

### DNA extraction and amplicon sequencing

Flies of each thermal stress treatment (non-stressed and heat-stressed flies) and sex were pooled into groups of 5 flies each: 10 pools of non-stressed females, 10 pools of non-stressed males, 10 pools of heat-stressed females, and 10 pools of heat-stressed males. To eliminate superficial bacteria, each pool was washed 3 washes with 0.5% hypochlorite sodium solution per 2 min each wash, 3 washes with 70% ethanol solution per 2 min each wash, and 3 washes with autoclaved water per 2 min each wash. Then, each pool was transferred in a Petri dish with sterile 1X PBS solution, where intestines of flies were removed and transferred to Eppendorf tubes with ice-cold sterile 1X PBS solution.

Genomic DNA was extracted from pooled guts using the GeneJet kit (Thermo-Fisher) following the protocol for negative-Gram and positive-Gram bacteria. Then, the V3-V4 hypervariable region of the 16S rRNA gene was amplified using a dual-indexing approach according to Fadrosh *et al.* (2014). Amplicon PCR was performed using modified 341F and 805F primers, which contained: 1) a linker sequence to bind amplicons to the Nextera XT DNA indexes; 2) a 12 bp barcode sequence to multiplex samples; 3) a 0 to 5 bp “heterogeneity spacer” to increase the heterogeneity of amplicon sequences; and 4) 16S rRNA gene universal primers (Table S1). The amplicon PCR mix had a final volume of 12.5 μl: 6.5 μl ultrapure water; 5 μl 2X Hot Start PCR Master Mix (Invitrogen); 0.25 μl 1 μM forward primer; 0.25 μl 1 μM reverse primer; and 0.5 μl template genomic DNA. Amplicon PCR cycle conditions were set up as follows: denaturation at 94 °C for 3 min, 35 amplification cycles at 94 °C for 45 s, 52 °C for 1 min, 72 °C for 70 s, and a final extension at 72 °C for 10 min. Amplified reactions were purified using an enzyme mix (exonuclease I and Fast AP, Invitrogen) to eliminate free primers and dNTPs, and then loaded on a 2% agarose gel stained with Sybr Safe (Invitrogen) to visualize the PCR products.

PCR product was quantified by fluorescence using the Quan-iT PicoGreen dsDNA kit (Invitrogen) and then, all samples were standardized at the lowest DNA concentration samples (7.78 ng/μl). The primer design allowed to multiplex 23 samples into two different sets of Nextera XT DNA indexes (Illumina Corporation, San Diego, CA). The index PCR had a final volume of 50 μl: 5μl amplicon PCR, 5 μl indexes (N701 and S502 for library 1, and N707 and S506 for library 2), 25 μl 2X Kapa Taq HotStart DNA Polymerase (Invitrogen), and 10 μl ultrapure water. Index PCR cycle conditions were set up as follows: denaturation at 94 °C for 3 min, 8 amplification cycles at 95 °C for 30 s, 55 °C for 30 s, 72 °C for 30 s, and a final extension at 72 °C for 5 min. PCR products were cleaning using the AMPure XT Bead kit (Beckman Coulter, Brea, CA) and quantified using the Qubit Fluorometer and Qubit dsDNA HS assay kit. Library 1 had a concentration of 37.2 ng/μl and library 2 had a concentration of 46.2 ng/μl, and both libraries were diluted at a concentration of 4 nM. Libraries were sequenced using an Illumina MiSeq sequencer (Illumina, San Diego, CA) and the MiSeq Reagent v3 (600 cycles). Sequencing was performed at AUSTRAL-omics Sequencing Core Facility at Universidad Austral de Chile.

### Metabarcoding analysis

After sequencing, 3,061,220 sequences were obtained. Raw sequence quality was inspected using FastQC (Andrews, 2010) and then filtered for a Q-value higher than 28 and sequences longer than 150 bp using the script Reads_Quality_Length_distribution.pl (Bálint et al., 2014). Forward and reverse filtered sequences were paired using PANDASeq with a minimum overlap of 5 bp (Masella et al., 2012). Paired-end sequences were trimmed to remove forward/reverse barcodes, heterogeneity spacers, and 16S rRNA gene primers. Quality-filtered and trimmed sequences were analyzed using QIIME v1.9.1 (Caporaso et al., 2010a). An open-reference OTU-picking strategy was used to generate operational taxonomic units (OTUs) using *usearch* v6.1 algorithm to cluster OTUs at 97 % of nucleotide identity. Taxonomy assignment was performed using *uclust* method (Edgar, 2010) against the Greengenes 16S rRNA gene database at 97% pairwise identity (version 13.8; Mcdonald *et al.*, 2012) as database reference. Finally, representative OTU sequences we aligned using PyNast and used to build a phylogenetic tree using FastTree. After this procedure, we retained 1,559,937 sequences assigned to 1,263 OTUs. After this, we performed two filtering steps: (1) remove mitochondrial-, chloroplast-, *Spiroplasma*-, and *Wolbachia*-related sequences; and (2) remove OTUs comprising less than 100 sequences. Retained sequences (total = 1,538,400 sequences : range = 746 – 59,210 sequences) and OTU number (total = 135) by sample are reported in Table S2. For diversity analyses, samples were rarified at 12,000 sequences according to the rarefaction curve (Fig. S1), which resulted in the removal of 4 samples (1FDRD, 2FDRD, 1FBRB, and 2FBRB; see Table S2).

### Statistical analyses

Assumptions of normality and homoscedasticity were checked for knockdown time. Then, knockdown time was analyzed using a two-factor analysis of variance (ANOVA) with microbiota condition (axenic and conventional flies) and sex (females and males) as main effects. Knockdown time at 35 and 36 °C showed a normal distribution, but knockdown time at 37 and 38 °C were squared-root transformed. We also tested the difference between survival curves of axenic and conventional flies at each static assays using the G-rho family test (long-rank test) using the *survival* R package (Therneau, 2020), and survival curves were plotted using the *survminer* R package (Kassambara et al., 2020).

We also analyzed the effects of heat stress, sex, and its interaction on the bacterial abundance, diversity indexes, and community structure of the gut microbiota. First, relative abundance at phylum and family taxonomical levels were obtained using *phyloseq* (Mcmurdie and Holmes, 2013) and *microbiome* (Lahti et al., 2017) packages for R, and then relative abundances were compared using a generalized linear model (GLM) assuming a quasibinomial distribution. Second, we analyzed the OTU relative abundances between non-stressed and heat-stressed flies for each sex by separated using the package *DESeq2* for R (Love et al., 2014). DESeq2 uses a negative-binomial model for count data, taking into account the zero-skewed distribution of the microbiome dataset. Significant differences between groups in OTU relative abundance were considered when the adjusted-*P* value (False Discovery Rate correction, FDR) was lower than 0.05. Third, OTU richness and Shannon diversity were estimated using the *microbiome* package for R (Lahti et al., 2017), whereas the phylogenetic diversity was estimated using QIIME. Diversity indexes were analyzed using a two-way ANOVA and *a posteriori* comparisons were performed using a Bonferroni t-test. Finally, we estimated the weighted-UniFrac distances among samples using QIIME, which was used as input to compare the bacterial community structure between thermal stress treatment (non-stressed and heat-stressed flies) and sexes (females and males). Bacterial community analysis was performed through a permutational multivariate analysis of variance (PERMANOVA) using *vegan* package for R (Oksanen et al., 2020)

All statistical analyses were performed using R version 4.0.3 (R Core Team, 2020) and RStudio version 1.3.959 (RStudio Team, 2020), and plots were made using *ggpubr* (Kassambara, 2020a) and *rstatix* (Kassambara, 2020b) packages for R.

## Results

### Gut microbiota and heat tolerance

As expected, knockdown time decreased when exposure temperature increased: the higher the exposure temperature, the shorter the knockdown time (Fig. 1). Evaluating the effect of the microbiota condition on heat tolerance at each static assay, we found that axenic flies exhibited a significantly lower heat tolerance than conventional flies in the assays at 35 °C (Table 1; Fig. 1A). For heat tolerance assayed at 36 °C, we found that ANOVA showed a marginally significant difference between axenic and conventional flies (Table 1), but the log-rank test showed significant differences between survival curves of axenic and conventional flies (Fig. 1B). On the other hand, thermal tolerance between axenic and conventional flies did not differ at 37 and 38 °C (Table 1; Fig. 1C and 1D, respectively). On the other hand, we did not find differences in heat tolerance between sexes at any thermal assay (Table 1). Finally, we only detected a significant interaction between the microbiota condition and sex at 35 °C (Table 1).

**Figure 1.**
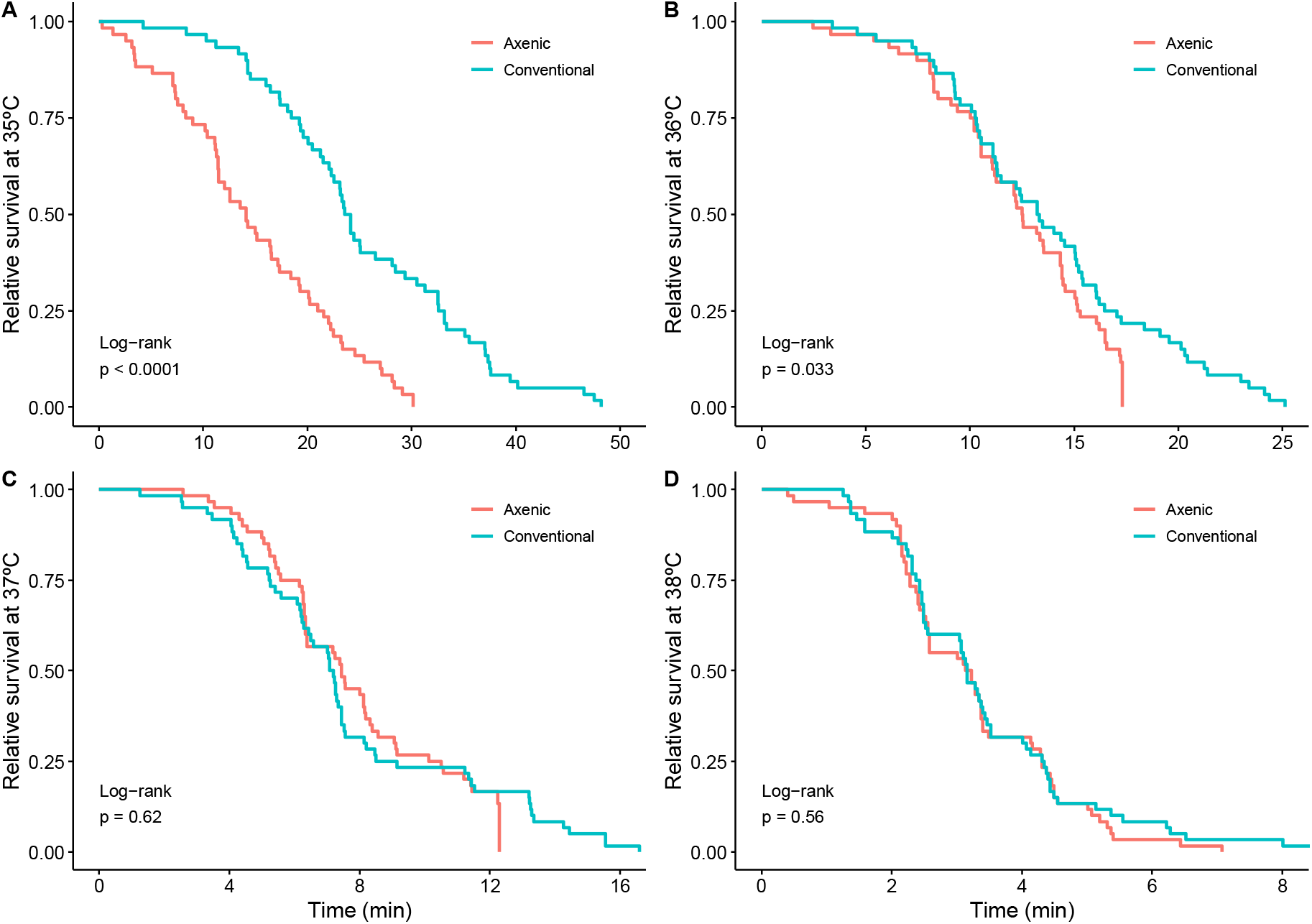
Relative survival (measured as knockdown time) of axenic (red) and conventional (green) flies estimated at different static assays (A: 35 °C, B: 36 °C, C: 37 °C, and D: 38 °C) in *Drosophila subobscura*. Survival curves (lines) were estimated using a Kaplan-Meier model, and shaded areas indicate 95% confidence interval of each survival curve.

**Table 1.** Results of the analysis of variance about the effects of microbiota condition (axenic and conventional flies), sex (female and male flies), and its interaction on heat tolerance (measures as the knockdown time in static assays) of *Drosophila subobscura*.

| Trait | Microbiota condition | Sex | Interaction |
| --- | --- | --- | --- |
| Knockdown time at 35°C | $F_{1,116} = 43.43$<br>$P = 1.35 \times 10^{-9}$ | $F_{1,116} = 1.54$<br>$P = 0.22$ | $F_{1,116} = 4.15$<br>$P = 0.04$ |
| Knockdown time at 36°C | $F_{1,116} = 3.47$<br>$P = 0.07$ | $F_{1,116} = 0.004$<br>$P = 0.95$ | $F_{1,116} = 1.16$<br>$P = 0.28$ |
| Knockdown time at 37°C | $F_{1,116} = 0.24$<br>$P = 0.63$ | $F_{1,116} = 3.18$<br>$P = 0.08$ | $F_{1,116} = 1.92$<br>$P = 0.17$ |
| Knockdown time at 38°C | $F_{1,116} = 0.31$<br>$P = 0.58$ | $F_{1,116} = 0.45$<br>$P = 0.50$ | $F_{1,116} = 1.63$<br>$P = 0.20$ |

### Gut microbiota composition

Gut microbiota of *D. subobscura* was dominated by bacteria belonging Actinobacteria (mean relative frequency ± SE = 0.09 ± 0.03), Bacteroidetes (mean relative frequency ± SE = 0.0002 ± 0.00004), Firmicutes (mean relative frequency ± SE = 0.66 ± 0.04), and Proteobacteria (mean relative frequency ± SE = 0.24 ± 0.04). In general, we found that the relative abundance of these phyla depended on heat stress and fly sex (Fig. 2). Actinobacteria increased their abundances from 0.2% in non-stressed females to 38.5% in heat-stressed females (GLM: *t* = −16.14, *P* = 9.6×10^−12^; Fig. S2A); whereas the increase was more moderated in male flies (GLM: *t* = −2.49, *P* = 0.02; Fig. S2A). Bacteroidetes relative abundance showed a significant interaction between heat stress and sex (GLM: *t* = 3.23, *P* = 0.003), with males showing an important reduction of Bacteroidetes abundance between non-stressed and heat-stressed flies (GLM: *t* = 5.58, *P* = 3.3×10^−5^; Fig. S2B) in comparison to female flies (GLM: *t* = 2.26, *P* = 0.04; Fig. S2B). Similarly, Proteobacteria abundance analysis showed a significant interaction between heat stress and sex (GLM: *t* = 7.77, *P* = 4.8×10^−9^; Fig. S2C): females showing similar abundance between non-stressed and heat-stressed flies (GLM: *t* = −1.69, *P* = 0.11), whereas heat stress induced an important reduction of Proteobacteria abundance in males (GLM: *t* = 7.92, *P* = 4.2×10^−5^). On the other hand, Firmicutes abundance showed an interaction response between heat stress and sex (GLM: *t* = −14.38, *P* = 5.2×10^−16^; Fig. S2D): females exposed to heat stress displayed a decrease in Firmicutes abundance (GLM: *t* = 29.18, *P* = 5.8×10^−16^), whereas heat stress induced an increase of Firmicutes abundance from 38.2% in non-stressed males to 89.4% in heat-stressed males (GLM: *t* = −7.99, *P* = 3.7×10^−7^).

**Figure 2.**
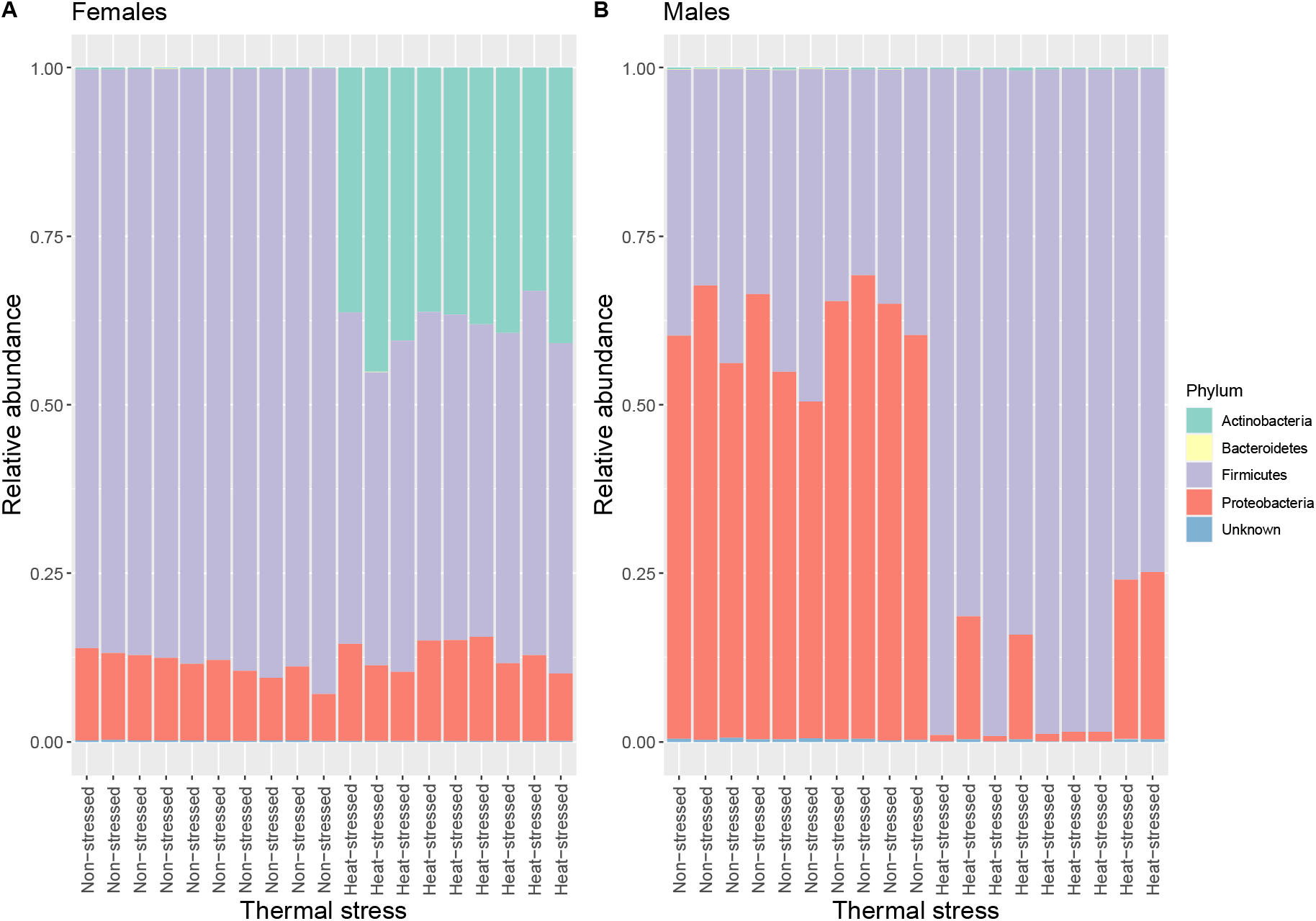
Relative abundance of bacterial composition (phylum) of the gut microbiota of non-stressed and heat-stressed female (A) and male (B) of *Drosophila subobscura*. Each color represents a bacterial phylum.

Among the most abundant bacterial families (abundance higher than 1%) associated with the gut of *D. subobscura*, we found Acetobacteraceae (mean relative frequency ± SE = 0.19 ± 0.03), Dermabacteraceae (mean relative frequency ± SE = 0.09 ± 0.03), Halomonadaceae (mean relative frequency ± SE = 0.02 ± 0.003), Lactobacillaceae (mean relative frequency ± SE = 0.45 ± 0.04), and Leuconostocaceae (mean relative frequency ± SE = 0.21 ± 0.05). Particularly, we focused on the relative abundances of acetic-acid and lactic-acid bacteria (AAB and LAB, respectively). We found a significant interaction between heat stress and sex for AAB (Acetobacteraceae) abundance (GLM: t = 6.97, *P* = 4.8×10^−8^; Fig. S3A), which is explained because whereas AAB abundance increased in heat-stressed females compared to non-stressed females (GLM: t = −7.20, *P* = 1.5×10^−6^) non-stressed males showed a higher abundance than heat-stressed males (GLM: t = 6.05, *P* = 1.3×10^−5^). On the other hand, LAB families exhibited contrasting responses to heat stress. Lactobacillaceae abundance showed a significant interaction between heat stress and sex (GLM: t = −2.80, *P* = 0.008; Fig. S3B): heat-stressed females a lower abundance of Lactobacillaceae than non-stressed females (GLM: t = 17.93, *P* = 1.8×10^−12^); whereas non-stressed and heat-stressed males showed similar Lactobacillaceae abundances (GLM: t = 1.67, *P* = 0.11). On the other hand, Leuconostocaceae abundance also showed a significant interaction between heat stress and sex (GLM: t = −5.45, *P* = 4.4×10^−6^; Fig. S3C): non-stressed females showed a higher Leuconostocaceae abundance than heat-stressed females (GLM: t = 2.37, *P* = 0.03); whereas heat-stressed males harbored a higher Leuconostocaceae abundance than non-stressed males (GLM: t = −4.85, *P* = 0.0001).

Analysis of OTU abundance (total = 135) showed that 42 OTUs significantly decreased their abundance in heat-stressed flies (blue circles in Fig. 3), whereas 51 OTUs significantly increased their abundance in heat-stressed flies (red circles in Fig. 3), and 42 OTUs did not change their abundance between non-stressed and heat-stressed flies (black circles in Fig. 3).

**Figure 3.**
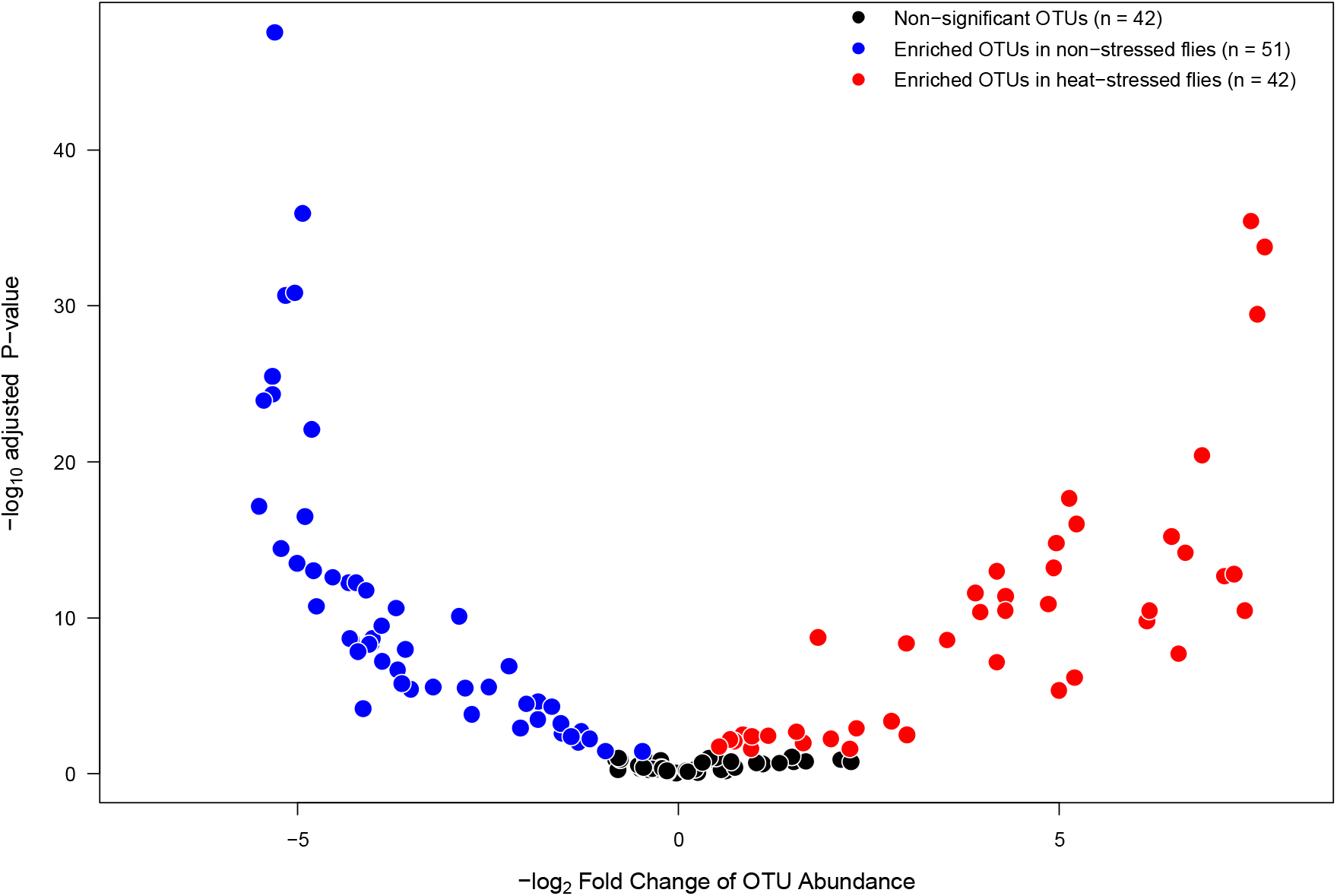
Volcano plot illustrating bacterial OTUs that showed significantly higher abundance in non-stressed flies (blue circles) or in heat-stressed flies (red circles) of *Drosophila subobscura*. OTUs that exhibit similar abundances in non-stressed and heat-stressed flies are represented by black circles.

### Gut microbiota diversity

OTU richness (Table S2) was significantly lower in heat-stressed flies than in non-stressed flies and not different between sexes, but we found a significant interaction between heat stress and sex (Table 2; Fig. 4A): thermal stress reduced the OTU number by 13.3% for female flies (Bonferroni t-test: *P* = 0.002), whereas this reduction was in 39.4% for male flies (Bonferroni t-test: *P* = 6.2×10^−7^). On the other hand, Shannon diversity (Table S2) showed non-significant individual effects associated with heat stress and sex, but a significant interaction was found (Table 2; Fig. 4B): non-stressed females harbor a lower diversity than heat-stressed females (Bonferroni t-test: *P* = 2.4×10^−10^), but non-stressed males showed higher diversity than heat-stressed males (Bonferroni t-test: *P* = 0.005). Finally, phylogenetic diversity (Table S2) was significantly lower in heat-stressed flies than in non-stressed flies and not different between sexes, but we found a significant interaction between heat stress and sex (Table 2; Fig. 4C): non-stressed and heat-stressed females showed a similar phylogenetic diversity (Bonferroni t-test: *P* = 0.18); whereas heat-stressed males harbor a lower phylogenetic diversity than non-stressed females (Bonferroni t-test: *P* = 5.5×10^−10^). Effects of heat stress and sex on the gut microbiota structure of *D. subobscura* showed a significant effect of heat stress (F_1,32_ = 73.35, *P* = 0.001, R^2^ = 0.19), sex (F_1,32_ = 33.48, *P* = 0.001, R^2^ = 0.09), and a significant interaction between heat stress and sex (F_1,32_ = 238.96, *P* = 0.001, R^2^ = 0.63). Heat stress had effects the gut microbiota structure of female and male flies, with each group clustering separately (Fig. 4D).

**Table 2.** Results of the analysis of variance about the effects of heat stress (non-stressed and heat-stressed flies), sex (female and male flies), and its interaction on bacterial diversity indices associated with the gut microbiota of *Drosophila subobscura*.

| Diversity index | Heat stress | Sex | Interaction |
| --- | --- | --- | --- |
| OTU number (richness) | $F_{1,32} = 124.81$<br>$P = 1.41 \times 10^{-12}$ | $F_{1,32} = 3.11$<br>$P = 0.09$ | $F_{1,32} = 33.43$<br>$P = 2.04 \times 10^{-6}$ |
| Shannon's diversity ($H'$) | $F_{1,32} = 0.41$<br>$P = 0.53$ | $F_{1,32} = 3.43$<br>$P = 0.07$ | $F_{1,32} = 63.19$<br>$P = 4.50 \times 10^{-9}$ |
| Phylogenetic diversity<br>(Faith's index) | $F_{1,32} = 93.37$<br>$P = 5.21 \times 10^{-11}$ | $F_{1,32} = 1.0$<br>$P = 0.33$ | $F_{1,32} = 43.50$<br>$P = 1.96 \times 10^{-7}$ |

**Figure 4.**
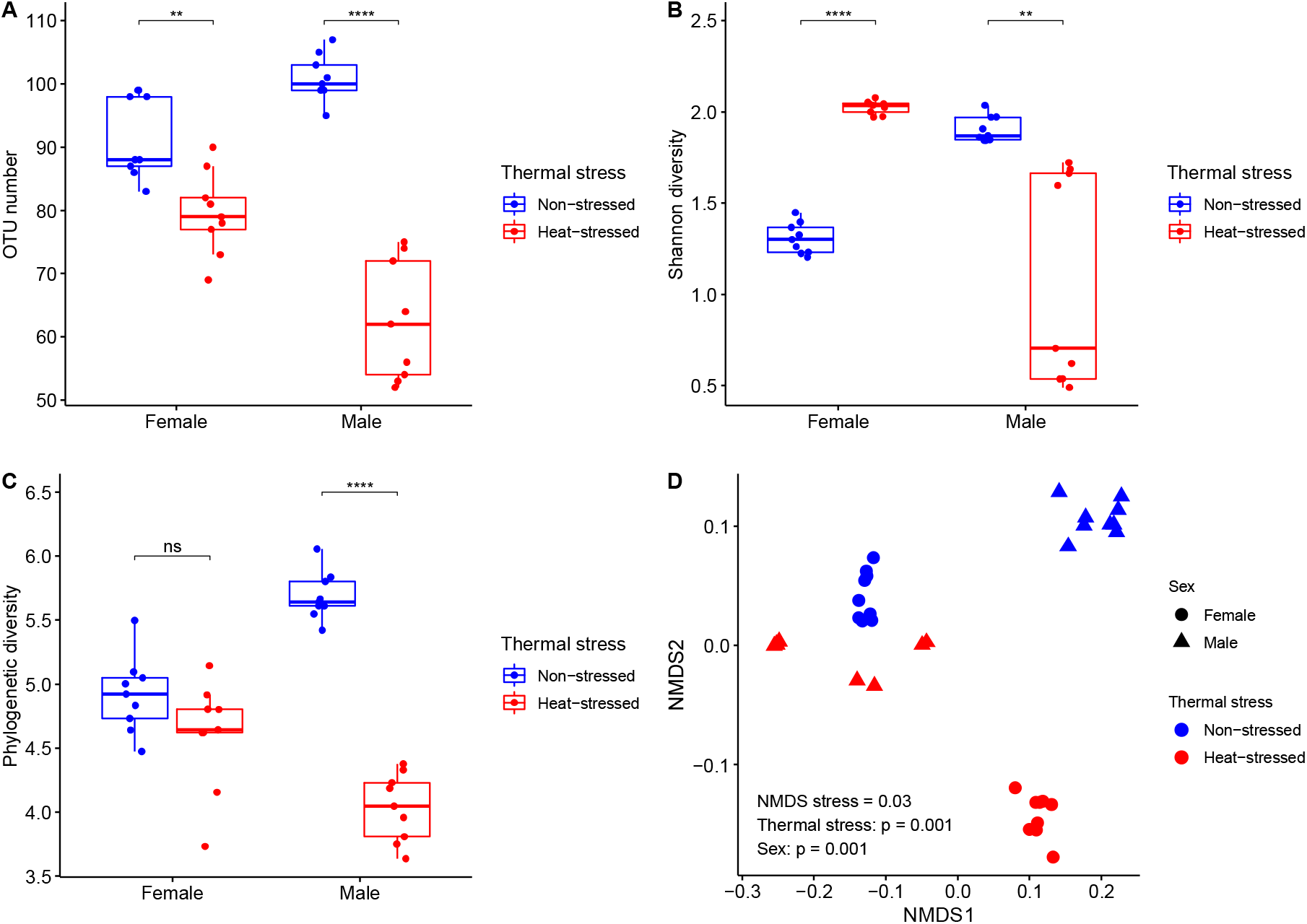
Bacterial diversity found in guts of non-stressed (blue) and heat-stressed (red) flies of *Drosophila subobscura* (females: circles, and males: triangles): (A) richness (OTU number); (B) Shannon’s diversity; (C) phylogenetic diversity (Faith’s index); and (D) bacterial community structure for gut microbiota using a non-metric multidimensional scaling (NMDS), which is based on weighted-UniFrac distances among samples. Box plots show median and interquartile range (IQR) and whiskers represent the 1.5*IQR. Symbols above boxplots denote non-significant (ns) or significant differences between non-stressed (blue) and heat-stressed flies (red) obtained from generalized-linear models (*: *P* < 0.05; **: *P* < 0.01; ***: *P* < 0.001; ****: *P* < 0.0001).

## Discussion

Global warming impacts the animal’s fitness, leading to an increased extinction risk in ectotherm species (Deutsch et al., 2008; Huey et al., 2012). Gut microbiota can contribute to host physiology leading to an increase of resistance to abiotic stressful conditions (Ferguson, 2018; Henry and Colinet, 2018). In the present work, we have studied the association between the gut microbiota and thermal physiology of *D. subobscura*, representing the first characterization of the gut microbiota for this species. Our findings provide evidence that gut microbiota influences heat tolerance, and that heat stress modifies the gut microbiota at taxonomical and diversity levels. These results demonstrate the sensitivity of the gut microbiota to transient heat stress, which can have negative impacts on host fitness.

### Gut microbiota and heat tolerance

Several studies have evaluated the role of the gut microbiota on cold and heat tolerance in ectotherms, finding different results. For instance, Henry and Colinet (2018) found that the gut microbiota contributes to cold tolerance, but they found no differences in heat tolerance between axenic and conventional flies of *D. melanogaster*. In the same line, Raza et al. (2020) found that the gut microbiota increases the tolerance to low temperatures in the dipteran *Bactrocera dorsalis*. On the other hand, a recent study found that the composition of the gut microbiota influences the heat tolerance of the western fence lizard (*Scleroporus occidentalis*), with a positive association between the abundance of genus *Anaerotignum* (Firmicutes) and heat tolerance (Moeller et al., 2020). The different results about the role of the gut microbiota on heat tolerance could be due the fact that heat tolerance depends on the methodology employed to measure it (Chown et al., 2009; Rezende et al., 2011; Castañeda et al., 2015), which can blurry physiological effects of different experimental treatments. To have a better approach to the thermal tolerance landscape (Rezende et al., 2014), we measured the heat tolerance at different temperatures from 35 °C (chronic thermal stress) to 38 °C (acute thermal stress). Interestingly, we only found a significant effect of the gut microbiota on heat tolerance when it was assayed at the lowest experimental temperature (35 °C): axenic flies tolerate this temperature for an average of 14.9 min, whereas conventional flies withstand for 25.5 min. A plausible explanation for these findings is the impact of gut microbiota on the host nutritional status (Ridley et al., 2012; Douglas, 2018a), which in turn determines the heat tolerance in ectotherms (Moghadam et al., 2018; Henry et al., 2019; Moeller et al., 2020). If this is true, thermal assays at high experimental temperatures (37 or 38 °C) occur so fast (7 and 3 min, respectively) that they cannot detect differences between axenic and conventional flies. Whereas, if gut microbiota impacts on nutritional status, conventional flies can withstand longer heat stress that axenic flies, but this difference can be only detected when there is enough time for energy reserves to be used in costly, resistance mechanisms such as heat tolerance (Feder and Hofmann, 1999; Calabria et al., 2012). Among these mechanisms, there is evidence that gut microbiota can influence the expression of heat-shock proteins in the gut epithelium (Liu et al., 2014; Arnal and Lalle, 2016), which represents a key response to mitigate cellular damage during thermal stress (Sørensen et al., 2003; Calabria et al., 2012).

### Gut microbiota composition

Recent studies have provided clear evidence about the impact of temperature on the gut microbiota of ectotherms (see Sepulveda and Moeller (2020) for a review). In general, these studies have used thermal acclimation (i.e. > 2 weeks) to evaluate changes in the gut microbiota composition and they have found that in warm temperatures, vertebrate ectotherms show a progressive decrease of bacteria belonging Firmicutes (Bestion et al., 2017; Fontaine et al., 2018), whereas warm-temperature acclimation led to an increase of the relative abundance of Proteobacteria invertebrate ectotherms (Berg et al., 2016; Moghadam et al., 2018; Horváthová et al., 2019). Here, we studied the effect of transient heat stress on the gut microbiota of *D. subobscura*, but we found a very different response of the bacterial composition when flies were exposed at 34 °C for 1 h. We found that the impact of heat stress led to an increase in the abundance in 37.8% of total OTUs, whereas 31.1% of OTUs decreased their abundance after heat stress. Interestingly, this short exposure to heat stress changed the gut microbiota composition differentially for each sex: heat stress induced a reduction in Firmicutes relative abundance, and an increase in Actinobacteria abundance; whereas for males, we observed an increase of Firmicutes and a decline of Proteobacteria abundances. Our findings suggest that temperature-induced changes in the gut microbiota of ectotherms can occur as fast as hours (present work), days (Sun et al., 2017), or weeks (Moghadam et al., 2018), which can explain the difference between our results and the expected increase of Proteobacteria in warm-acclimated ectotherms. Additionally, this difference can be explained by the fact that we analyzed the impact of temperature on the gut microbiota in both sexes, whereas other studies have assessed this impact only using only males (Moghadam *et al.*, 2018 and Horváthová *et al.*, 2019).

At the family level, we found that the gut microbiota of *D. subobscura* was dominated by acetic-acid (*Acetobacteraceae*) and lactic-acid (*Lactobacillaceae* and *Leuconostoceae*) bacteria, which is a common characteristic in *Drosophila* species (Douglas, 2018b). Regarding the effect of temperature on the bacterial family composition, *D. melanogaster* acclimated in warm conditions show a higher abundance of *Acetobacter* bacteria (AAB) and lower abundance of *Leuconostoc* bacteria (LAB) in comparison to cold-acclimated flies (Moghadam et al., 2018). Interestingly, this temperature-induced response of the gut microbiota composition under laboratory conditions matches wild populations, where low-latitude populations of *D. melanogaster* showed a higher AAB abundance and lower LAB abundance compared to high-latitude populations (Walters et al., 2020). Here, we found that AAB and LAB abundance changed with thermal stress, but these changes depended on the fly sex. In general, thermal stress reduced the *Acetobacteraceae* (AAB) and *Lactobacillaceae* (LAB) abundances, but the relative abundance of *Leuconostoceae* (LAB) increased in heat-stressed flies. Differences between *D. melanogaster* and *D. subobscura* can be explained: (1) because they have traditionally fed on different diets, which is known to impact the gut microbiota composition (Jehrke et al., 2018; Obadia et al., 2018); or (2) just because they diverged around 40 million years ago (Gibbs and Matzkin, 2001), resulting in different evolutionary histories under different environmental contexts. Therefore, comparative studies are needed to understand the thermal plasticity of the gut microbiota in a wider range of *Drosophila* species.

### Gut microbiota diversity

Temperature also has important effects on the diversity and structure of the gut microbiota in ectotherms (Sepulveda and Moeller, 2020). In general, warm conditions lead to a decrease of OTU number (richness) and diversity of gut microbiota in ectotherms (Bestion et al., 2017; Kokou et al., 2018; Li et al., 2018). Here, we found that heat stress induces a reduction in OTU number and phylogenetic diversity decreased in heat-stressed flies, being these effects more important in males than females. Similar response of richness and phylogenetic diversity is not surprising because commonly both diversity indices are highly correlated (Tôrres and Diniz-Filho, 2004). Conversely, Shannon diversity increased in heat-stressed females but decreased in heat-stressed males, which can be associated with changes in the abundance of some phyla in response to heat tolerance. Indeed, non-stressed females and heat-stressed males showed a more similar community structure compared to the other groups. This can be explained because the gut bacterial community of the non-stressed females was dominated by Firmicutes (88.4%) and Proteobacteria (11.2%), which was relatively similar for heat-stressed males: Firmicutes (89.7%) and Proteobacteria (9.8%). Changes in the gut microbiota diversity are common in ectotherms exposed to warm conditions and it could be explained because some bacterial species have higher thermal tolerance or they can tolerate better indirect effects of heat stress such as the production of reactive oxygen species by hosts as s response to heat stress (Lian et al., 2020). However, our study had some limitations to explain the proximal causes about changes in bacterial abundances and future steps should be focused to explore the resistance mechanisms in members of the gut microbiota.

## Conclusions

Temperature induces changes in the gut microbiota of ectotherms, regardless of how long organisms have been exposed to warm conditions. Here, we demonstrate that these changes are different for both sexes, and future studies should assess the sexual dimorphism in gut microbiota responses to abiotic and biotic factors. These changes in gut microbiota have consequences on the physiological mechanism as thermal resistance, which can impact on host fitness, population risk extinction, and vulnerability of ectotherms to current and future climatic conditions. Research about the role of gut microbiota on the adaptive response to climate change is a new venue and future research needs to balance mechanistic approaches to understand the host-microbiota interactions and holistic approaches to know the role of gut microbiota on the ecology and evolution of ectotherms.

## Author’s contributions

A.J. performed experiments, analyze the dataset and approved the final version of the manuscript. L.E.C. conceived the original idea, designed experiments, conducted bioinformatic and statistical analyses, provided funds for all experiments, and wrote the manuscript.

## Acknowledgments

We thank Andrea Silva from AUSTRAL-omics Sequencing Core Facility (Universidad Austral de Chile) for her advice and support during amplicon sequencing. A.J. thanks an internal scholarship of the Master Program in Genetics (Facultad de Ciencias, Universidad Austral de Chile).

## Data accessibility

Data available from Figshare: https://figshare.com/s/07258a71c3074fa59b1e. For copyright reasons, amplicon sequences will be available once the manuscript has been published.

## Supplementary Material

**Figure S1.**
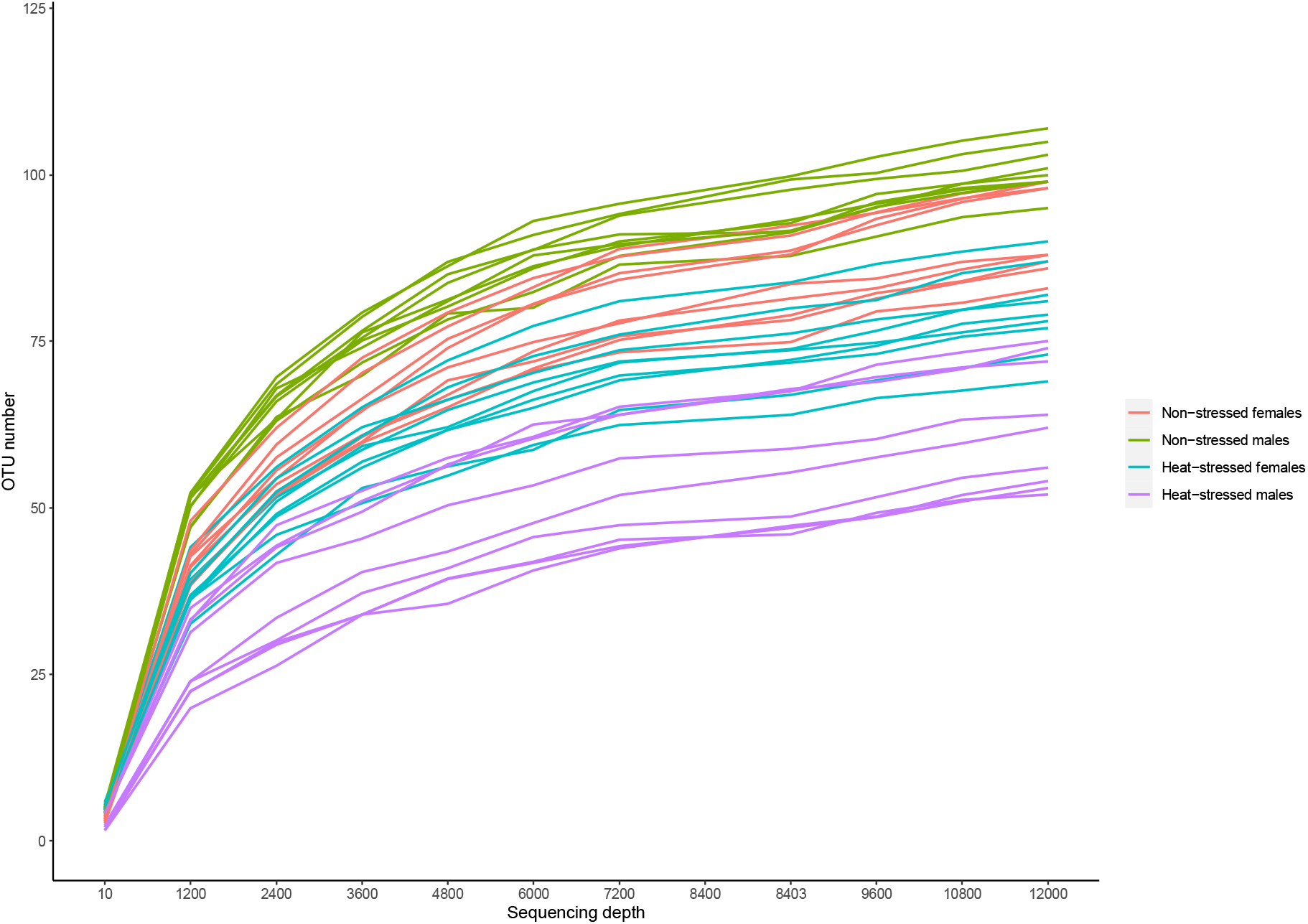
Rarefaction plots indicating OTU number and sequencing depth after rarefaction at 12,000 sequences. Samples were taken from non-stressed female (orange), non-stressed male (green), heat-stressed female (lightblue), and heat-stressed male (purples) flies of *D. subobscura*.

**Figure S2.**
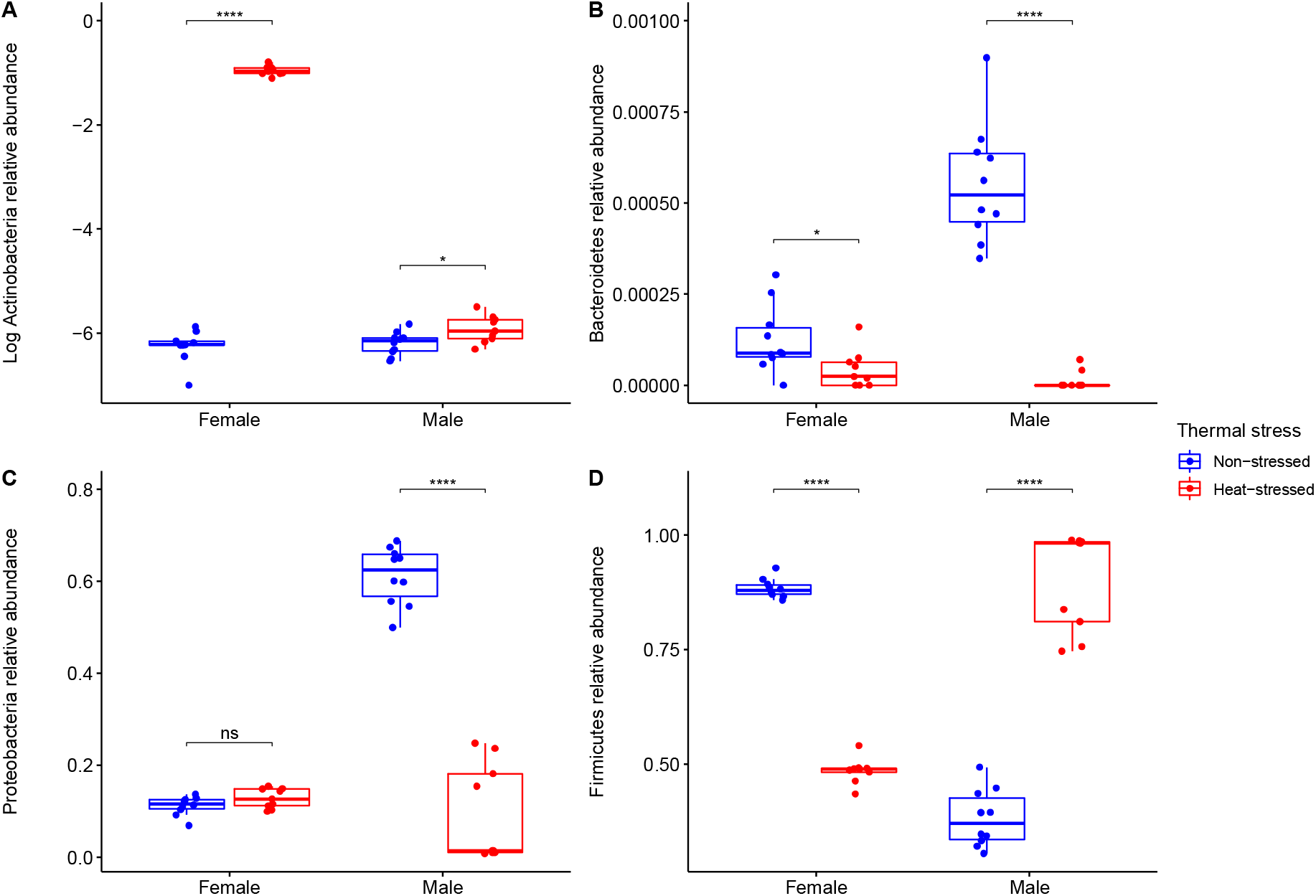
Relative abundances of the phyla (A) Actinobacteria, (B) Bacteroidetes, (C) Proteobacteria, and (D) Firmicutes associated with the guts of non-stressed (blue) and heat-stressed (red) flies (females and males) of *Drosophila subobscura*. Box plots show median and interquartile range (IQR) and whiskers represent the 1.5*IQR. Symbols above boxplots denote non-significant (ns) or significant differences between non-stressed (blue) and heat-stressed flies (red) obtained from generalized-linear models (*: *P* < 0.05; **: *P* < 0.01; ***: *P* < 0.001; ****: *P* < 0.0001).

**Figure S3.**
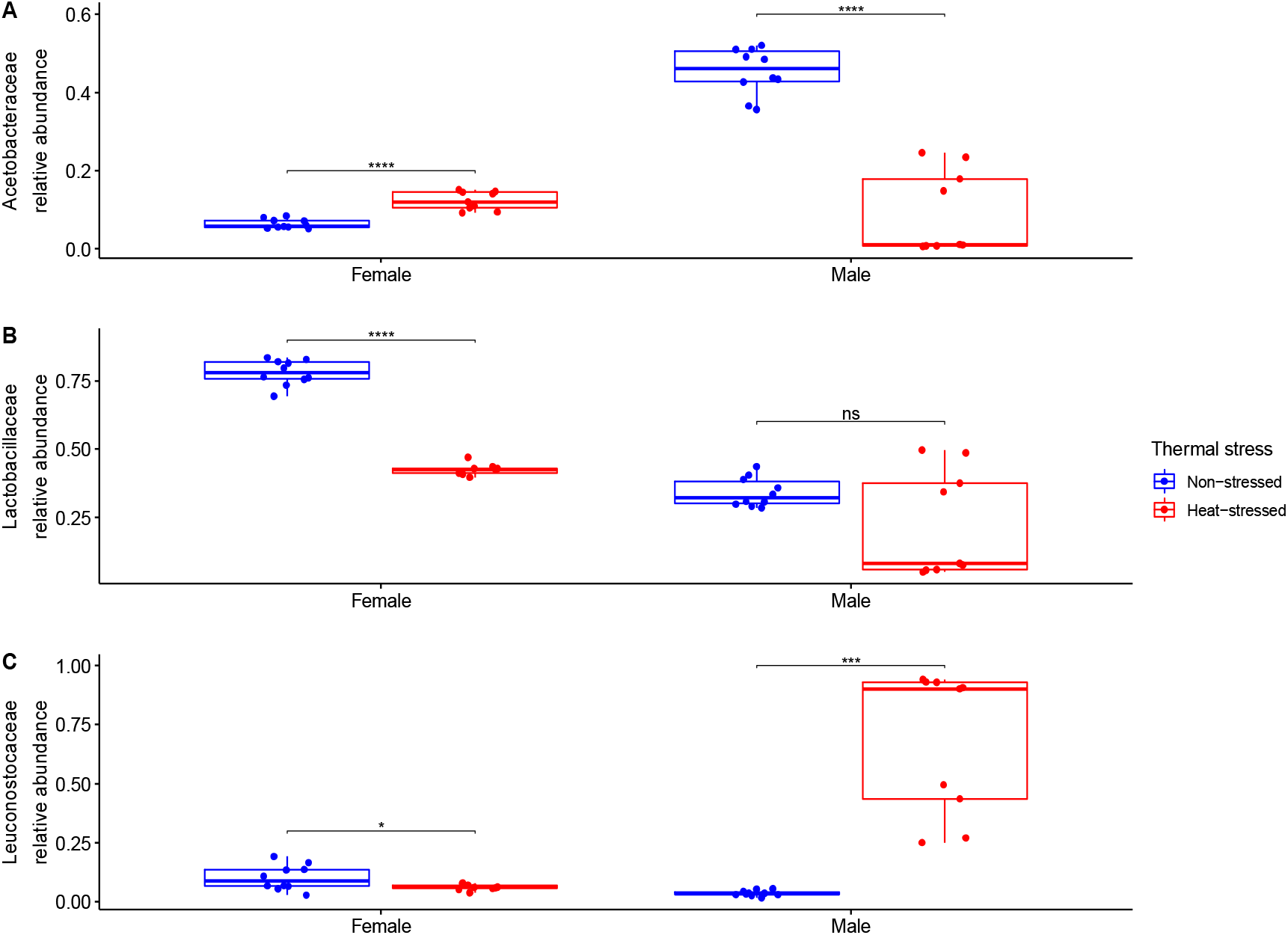
Relative abundances of the bacterial families (A) Acetobacteraceae, (B) Lactobacillaceae, and (C) Leuconostoceae associated with the guts of non-stressed (blue) and heat-stressed (red) flies (females and males) of *Drosophila subobscura*. Box plots show median and interquartile range (IQR) and whiskers represent the 1.5*IQR. Symbols above boxplots denote non-significant (ns) or significant differences between non-stressed (blue) and heat-stressed flies (red) obtained from generalized-linear models (*: *P* < 0.05; **: *P* < 0.01; ***: *P* < 0.001; ****: *P* < 0.0001).

**Table S1.**
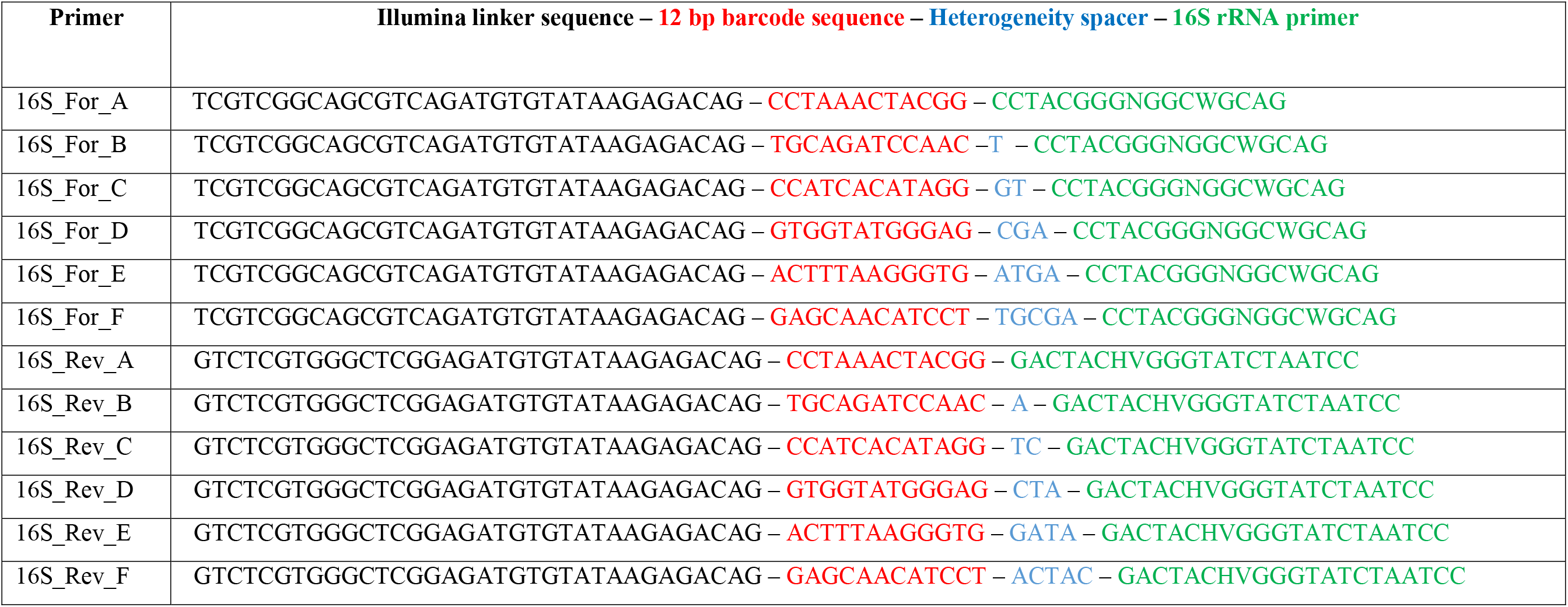
Modified 341F and 805F primers for bacterial 16S rRNA gene used in this study. Each primer contains: a linker sequence to bind amplicons to the Nextera XT DNA indexes (black letters); a 12 bp barcode sequence to multiplex samples (red letters); a 0 to 5 bp “heterogeneity spacer” (blue letters), and 16S rRNA gene universal primers (green letters).

**Table S2.** Information for each sample about the thermal stress and sex of Drosophila subobscura included in this study. Number of sequences, removal after rarefaction at 12,000 sequences, and richness (OTU number), Shannon diversity, and phylogenetic diversity estimates for each sample are provided. NA means non-available estimate because sample was removed after rarefaction (see Material and Methods section).

| Sample ID | Sex | Thermal stress | Sequences | Sample removed after rarefaction | Richness (OTU number) | Shannon diversity | Phylogenetic diversity |
| --- | --- | --- | --- | --- | --- | --- | --- |
| 1FDRD | Female | Heat-stressed | 746 | Yes | NA | NA | NA |
| 2FDRD | Male | Heat-stressed | 860 | Yes | NA | NA | NA |
| 2FBRB | Male | Non-stressed | 5748 | Yes | NA | NA | NA |
| 1FBRB | Female | Non-stressed | 6596 | Yes | NA | NA | NA |
| 2FCRC | Female | Heat-stressed | 12496 | No | 90 | 2.0783283 | 5.14382 |
| 1FCRC | Male | Heat-stressed | 13042 | No | 62 | 0.7040019 | 4.18603 |
| 1FARA | Male | Non-stressed | 15864 | No | 105 | 1.9065749 | 6.05509 |
| 2FARA | Female | Non-stressed | 17341 | No | 83 | 1.203535 | 4.47469 |
| 2FBRD | Male | Heat-stressed | 24885 | No | 72 | 1.5965272 | 4.23032 |
| 2FCRD | Male | Heat-stressed | 28514 | No | 74 | 1.6631817 | 4.33094 |
| 2FCRB | Male | Non-stressed | 33992 | No | 99 | 2.0364269 | 5.64249 |
| 2FCRA | Female | Non-stressed | 35376 | No | 88 | 1.26127 | 4.64091 |
| 2FDRC | Female | Heat-stressed | 38526 | No | 77 | 1.9712162 | 4.80308 |
| 2FBRC | Female | Heat-stressed | 39676 | No | 87 | 2.045393 | 4.62237 |
| 2FARC | Female | Heat-stressed | 40144 | No | 78 | 2.0240818 | 4.80176 |
| 1FCRA | Male | Non-stressed | 41559 | No | 103 | 1.8614536 | 5.54767 |
| 2FERC | Female | Heat-stressed | 41693 | No | 81 | 2.0365678 | 4.91643 |
| 2FDRB | Male | Non-stressed | 42681 | No | 99 | 1.8427923 | 5.61006 |
| 2FBRA | Female | Non-stressed | 44190 | No | 88 | 1.2241704 | 4.83322 |
| 2FARD | Male | Heat-stressed | 45337 | No | 75 | 1.7221588 | 4.37624 |
| 2FERB | Male | Non-stressed | 45609 | No | 99 | 1.8698738 | 5.42088 |
| 1FDRC | Male | Heat-stressed | 45911 | No | 56 | 0.6206609 | 3.63735 |
| 2FERA | Female | Non-stressed | 46086 | No | 99 | 1.3249281 | 5.09436 |
| 2FDRA | Female | Non-stressed | 46127 | No | 98 | 1.2309471 | 5.04824 |
| 1FERD | Female | Heat-stressed | 47339 | No | 82 | 2.0531454 | 4.62203 |
| 1FERB | Female | Non-stressed | 48321 | No | 86 | 1.4479118 | 4.73222 |
| 1FBRC | Male | Heat-stressed | 48350 | No | 53 | 0.5368444 | 3.75234 |
| 1FARC | Male | Heat-stressed | 49183 | No | 52 | 0.5356601 | 4.04646 |
| 1FERA | Male | Non-stressed | 49422 | No | 101 | 1.9705597 | 5.80055 |
| 1FCRD | Female | Heat-stressed | 49884 | No | 79 | 2.044334 | 4.64284 |
| 1FERC | Male | Heat-stressed | 50290 | No | 54 | 0.4892269 | 3.95857 |
| 1FBRD | Female | Heat-stressed | 50902 | No | 73 | 1.9746317 | 4.1562 |
| 1FDRB | Female | Non-stressed | 51134 | No | 98 | 1.366587 | 5.00307 |
| 2FARB | Male | Non-stressed | 51351 | No | 107 | 1.9728177 | 5.8345 |
| 1FCRB | Female | Non-stressed | 53051 | No | 99 | 1.3970142 | 5.49794 |
| 1FDRA | Male | Non-stressed | 53163 | No | 100 | 1.8445902 | 5.66431 |
| 1FARD | Female | Heat-stressed | 53901 | No | 69 | 2.0003884 | 3.73369 |
| 1FBRA | Male | Non-stressed | 54842 | No | 95 | 1.8485612 | 5.60939 |
| 2FERD | Male | Heat-stressed | 55058 | No | 64 | 1.6865227 | 3.8092 |
| 1FARB | Female | Non-stressed | 59210 | No | 87 | 1.3002729 | 4.92085 |

## References

Alberdi, A., Aizpurua, O., Bohmann, K., Zepeda-Mendoza, M. L., and Gilbert, M. T. P. (2016). Do vertebrate gut metagenomes confer rapid ecological adaptation? Trends Ecol. Evol. 31, 689–699. doi:10.1016/j.tree.2016.06.008.

Andrews, S. (2010). FastQC: a quality control tool for high throughput sequence data. Available at http://www.bioinformatics.babraham.ac.uk/projects/fastqc.

Angilletta, M. J. (2009). Thermal Adaptation: a Theoretical and Empirical Synthesis. Oxford University Press.

Arnal, M., and Lalle, J. (2016). Gut epithelial inducible heat-shock proteins and their modulation by diet and the microbiota. Nutr. Rev. 74, 181–197. doi:10.1093/nutrit/nuv104.

Bálint, M., Schmidt, P.-A., Sharma, R., Thines, M., and Schmitt, I. (2014). An Illumina metabarcoding pipeline for fungi. Ecol. Evol. 4, 2642–2653. doi:10.1002/ece3.1107.

Berg, M., Stenuit, B., Ho, J., Wang, A., Parke, C., Knight, M., et al. (2016). Assembly of the Caenorhabditis elegans gut microbiota from diverse soil microbial environments. ISME J. 10, 1998–2009. doi:10.1038/ismej.2015.253.

Bestion, E., Jacob, S., Zinger, L., Gesu, L. Di, Richard, M., White, J., et al. (2017). Climate warming reduces gut microbiota diversity in a vertebrate ectotherm. Nat. Ecol. Evol. 1, 0161. doi:10.1038/s41559-017-0161.

Broderick, N. A., and Lemaitre, B. (2012). Gut-associated microbes of Drosophila melanogaster. Gut Microbes 3, 307–321. doi:10.4161/gmic.19896.

Calabria, G., Dolgova, O., Rego, C., Castañeda, L. E., Rezende, E. L., Balanyà, J., et al. (2012). Hsp70 protein levels and thermotolerance in Drosophila subobscura: A reassessment of the thermal co-adaptation hypothesis. J. Evol. Biol. 25, 691–700. doi:10.1111/j.1420-9101.2012.02463.x.

Caporaso, J. G., Kuczynski, J., Stombaugh, J., Bittinger, K., Bushman, F. D., Costello, E. K., et al. (2010a). QIIME allows analysis of high- throughput community sequencing data. Nat. Methods 7, 335–336. doi:10.1038/nmeth0510-335.

Caporaso, J. G., Lauber, C. L., Walters, W. A., Berg-lyons, D., Lozupone, C. A., Turnbaugh, P. J., et al. (2010b). Global patterns of 16S rRNA diversity at a depth of millions of sequences per sample. Proc. Natl. Acad. Sci. U. S. A. 108, 4516–4522. doi:10.1073/pnas.1000080107/-/DCSupplemental.www.pnas.org/cgi/doi/10.1073/pnas.1000080107.

Castañeda, L. E., Rezende, E. L., and Santos, M. (2015). Heat tolerance in Drosophila subobscura along a latitudinal gradient: Contrasting patterns between plastic and genetic responses. Evolution (N. Y). 69, 2721–2734. doi:10.1111/evo.12757.

Castañeda, L. E., Soriano, V. R., Mesas, A., and Roff, D. A. (2019). Evolutionary potential of thermal preference and heat tolerance in Drosophila subobscura. J. Evol. Biol. 32, 818–824. doi:10.1111/jeb.13483.

Chown, S. L., Jumbam, K. R., Sørensen, J. G., and Terblanche, J. S. (2009). Phenotypic variance, plasticity and heritability estimates of critical thermal limits depend on methodological context. Funct. Ecol. 23, 133–140. doi:10.1111/j.1365-2435.2008.01481.x.

Deutsch, C. A., Tewksbury, J. J., Huey, R. B., Sheldon, K. S., Ghalambor, C. K., Haak, D. C., et al. (2008). Impacts of climate warming on terrestrial ectotherms across latitude. Proceeding Natl. Acad. Sci. 105, 6668–6672.

Douglas, A. E. (2018a). Fundamentals of Microbiome Science: How Microbes Shape Animal Biology. Princeton University Press.

Douglas, A. E. (2018b). The Drosophila model for microbiome research. Lab Anim. (NY). 47, 157–164. doi:10.1038/s41684-018-0065-0.

Dunbar, H. E., Wilson, A. C. C., Ferguson, N. R., and Moran, N. A. (2007). Aphid thermal tolerance is governed by a point mutation in bacterial symbionts. PLoS Biol. 5, e96. doi:10.1371/journal.pbio.0050096.

Edgar, R. C. (2010). Search and clustering orders of magnitude faster than BLAST. Bioinformatics 26, 2460–2461. doi:10.1093/bioinformatics/btq461.

Fadrosh, D. W., Ma, B., Gajer, P., Sengamalay, N., Ott, S., Brotman, R. M., et al. (2014). An improved dual-indexing approach for multiplexed 16S rRNA gene sequencing on the Illumina MiSeq platform. Microbiome 2, 6.

Feder, M. E., and Hofmann, G. E. (1999). Heat-shock proteins, molecular chaperones, and the stress response: evolutionary and ecological physiology. Annu. Rev. Physiol. 61, 243–282.

Ferguson, L. V (2018). Seasonal shifts in the insect gut microbiome are concurrent with changes in cold tolerance and immunity. Funct. Ecol. 32, 2357–2368.

Fontaine, S. S., Novarro, A. J., and Kohl, K. D. (2018). Environmental temperature alters the digestive performance and gut microbiota of a terrestrial amphibian. J. Exp. Biol. 221, 187559. doi:10.1242/jeb.187559.

Gibbs, a G., and Matzkin, L. M. (2001). Evolution of water balance in the genus Drosophila. J. Exp. Biol. 204, 2331–8. Available at: http://www.ncbi.nlm.nih.gov/pubmed/11507115.

Gruntenko, N. Е., Ilinsky, Y. Y., Adonyeva, N. V, Burdina, E. V, Bykov, R. A., Menshanov, P. N., et al. (2017). Various Wolbachia genotypes differently influence host Drosophila dopamine metabolism and survival under heat stress conditions. BMC Evol. Biol. 17, 252. doi:10.1186/s12862-017-1104-y.

Henry, Y., and Colinet, H. (2018). Microbiota disruption leads to reduced cold tolerance in Drosophila flies. Sci. Nat. 105, 59.

Henry, Y., Overgaard, J., and Colinet, H. (2019). Dietary nutrient balance shapes phenotypic traits of Drosophila melanogaster in interaction with gut microbiota. Comp. Biochem. Physiol. Part A 241, 110626. doi:10.1016/j.cbpa.2019.110626.

Hoffmann, A. a., Sørensen, J. G., and Loeschcke, V. (2003). Adaptation of Drosophila to temperature extremes: bringing together quantitative and molecular approaches. J. Therm. Biol. 28, 175–216. doi:10.1016/S0306-4565(02)00057-8.

Hoffmann, A. a, and Sgrò, C. M. (2011). Climate change and evolutionary adaptation. Nature 470, 479–85. doi:10.1038/nature09670.

Horváthová, T., Koz, J., and Bauchinger, U. (2019). Vanishing benefits: the loss of actinobacterial symbionts at elevated temperatures. J. Therm. Biol. 82, 222–228.

Hoye, B. J., and Fenton, A. (2018). Animal host: microbe interactions. J. Anim. Ecol. 87, 315–319. doi:10.1111/1365-2656.12788.

Huey, R. B., Kearney, M. R., Krockenberger, A., Holtum, J. A. M., Jess, M., and Williams, S. E. (2012). Predicting organismal vulnerability to climate warming: roles of behaviour, physiology and adaptation. Philos. Trans. R. Soc. Lond. B. Biol. Sci. 367, 1665–1679. doi:10.1098/rstb.2012.0005.

Jehrke, L., Stewart, F. A., Droste, A., and Beller, M. (2018). The impact of genome variation and diet on the metabolic phenotype and microbiome composition of Drosophila melanogaster. Sci. Rep. 8, 1–15. doi:10.1038/s41598-018-24542-5.

Kassambara, A. (2020a). ggpubr: “ggplot2” based publication ready plots. R package version 0.4.0. https://CRAN.R-project.org/package=ggpubr.

Kassambara, A. (2020b). rstatix: pipe-friendly framework for basic statistical tests. R package version 0.6.0. URL: https://CRAN.R-project.org/package=rstatix.

Kassambara, A., Kosinski, M., and Biecek, P. (2020). survminer: Drawing Survival Curves using “ggplot2”. R package version 0.4.8. URL: https://CRAN.R-project.org/package=survminer.

Kokou, F., Sasson, G., Nitzan, T., and Doron-faigenboim, A. (2018). Host genetic selection for cold tolerance shapes microbiome composition and modulates its response to temperature. Elife 7, e36398.

Koyle, M. L., Veloz, M., Judd, A. M., Wong, A. C.-N., Newell, P. D., Douglas, A. E., et al. (2016). Rearing the fruit fly Drosophila melanogaster under axenic and gnotobiotic conditions. J. Vis. Exp., e54219. doi:10.3791/54219.

Lahti, L., Shetty, S., Turaga, N., Obenchain, V., Salojärvi, J., Gilmore, R., et al. (2017). Tools for microbiome analysis in R. Version. Available at http://microbiome.github.com/microbiome.

Li, Y., Yang, N., Liang, X., Yoshida, A., Osatomi, K., and Grove, T. J. (2018). Elevated seawater temperatures secrease microbial diversity in the gut of Mytilus coruscus. Front. Physiol. 9, 839. doi:10.3389/fphys.2018.00839.

Lian, P., Braber, S., Garssen, J., Wichers, H. J., Folkerts, G., Fink-Gremmels, J., et al. (2020). Beyond heat stress: intestinal integrity disruption and mechanism-based intervention strategies. Nutrients 12, 734.

Liu, H., Dicksved, J., Lundh, T., and Lindberg, J. E. (2014). Heat shock proteins: intestinal gatekeepers that are influenced by dietary components and the gut microbiota. Pathogens, 187–210. doi:10.3390/pathogens3010187.

Love, M. I., Huber, W., and Anders, S. (2014). Moderated estimation of fold change and dispersion for RNA-seq data with DESeq2. Genome Biol. 15, 550.

Macke, E., Tasiemski, A., Massol, F., Callens, M., and Decaestecker, E. (2017). Life history and eco-evolutionary dynamics in light of the gut microbiota. Oikos 126, 508–531. doi:10.1111/oik.03900.

Masella, A. P., Bartram, A. K., Truszkowski, J. M., Brown, D. G., and Neufeld, J. D. (2012). PANDAseqC: PAired-eND Assembler for Illumina sequences. BMC Bioinformatics 13, 31.

Mcdonald, D., Price, M. N., Goodrich, J., Nawrocki, E. P., Desantis, T. Z., Probst, A., et al. (2012). An improved Greengenes taxonomy with explicit ranks for ecological and evolutionary analyses of bacteria and archaea. ISME J. 7, 610–618. doi:10.1038/ismej.2011.139.

Mcmurdie, P. J., and Holmes, S. (2013). phyloseq◻: an R rackage for reproducible interactive analysis and graphics of microbiome census data. PLoS One 8, e61217. doi:10.1371/journal.pone.0061217.

Moeller, A. H., Ivey, K., Cornwall, M. B., Herr, K., Rede, J., Taylor, E. N., et al. (2020). Lizard gut microbiome changes with temperature and is associated with heat tolerance. Appl. Environ. Microbiol. 18, e01181–20. doi:10.1128/AEM.01181-20.

Moghadam, N. N., Mai, P., Kristensen, T. N., Jonge, N. De, and Bahrndorff, S. (2018). Strong responses of Drosophila melanogaster microbiota to developmental temperature. Fly (Austin). 12, 1–12. doi:10.1080/19336934.2017.1394558.

Montllor, C. B., Maxmen, A., and Purcell, A. H. (2002). Facultative bacterial endosymbionts benefit pea aphids Acyrthosiphon pisum under heat stress. Ecol. Entomol. 27, 189–195.

Obadia, B., Keebaugh, E. S., Yamada, R., Ludington, W. B., and Ja, W. W. (2018). Diet influences host-microbiota associations in Drosophila. Proc. Natl. Acad. Sci. U. S. A. 115, 201804948. doi:10.1073/pnas.1804948115.

Oksanen, J., Blanchet, F. G., Friendly, M., Kindt, R., Legendre, P., McGlinn, D., et al. (2020). Vegan: community ecology package. R package version 2.5-7.

R Core Team (2020). R: A language and environment for statistical computing. R Foundation for Statistical Computing, Vienna, Austria. URL: https://www.R-project.org/.

Raza, M. F., Wang, Y., Cai, Z., Bai, S., Yao, Z., Awan, A., et al. (2020). Gut microbiota promotes host resistance to low-temperature stress by stimulating its arginine and proline metabolism pathway in adult Bactrocera dorsalis. PLoS Pathog. 16, e1008441. doi:10.1371/journal.ppat.1008441.

Renoz, F., Pons, I., and Hance, T. (2019). Evolutionary responses of mutualistic insect: bacterial symbioses in a world of fluctuating temperatures. Curr. Opin. Insect Sci. 35, 20–26. doi:10.1016/j.cois.2019.06.006.

Rezende, E. L., Castañeda, L. E., and Santos, M. (2014). Tolerance landscapes in thermal ecology. Funct. Ecol. 28. doi:10.1111/1365-2435.12268.

Rezende, E. L., Tejedo, M., and Santos, M. (2011). Estimating the adaptive potential of critical thermal limits: methodological problems and evolutionary implications. Funct. Ecol. 25, 111–121. doi:10.1111/j.1365-2435.2010.01778.x.

Ridley, E. V., Wong, A. C. N., Westmiller, S., and Douglas, A. E. (2012). Impact of the resident microbiota on the nutritional phenotype of Drosophila melanogaster. PLoS One 7, e36765. doi:10.1371/journal.pone.0036765.

Romano, M. (2017). Gut microbiota as a trigger of accelerated directional adaptive evolution: Acquisition of herbivory in the context of extracellular vesicles, microRNAs and inter-kingdom crosstalk. Front. Microbiol. 8, 1–7. doi:10.3389/fmicb.2017.00721.

RStudio Team (2020). RStudio: integrated development for R. RStudio, PBC, Boston, MA. URL http://www.rstudio.com/.

Russell, J. A., and Moran, N. A. (2006). Costs and benefits of symbiont infection in aphids: variation among symbionts and across temperatures. Proceeding R. Soc. London Ser. B 273, 603–610. doi:10.1098/rspb.2005.3348.

Sepulveda, J., and Moeller, A. H. (2020). The effects of temperature on animal gut microbiomes. Front. Microbiol. 11, 384. doi:10.3389/fmicb.2020.00384.

Sørensen, J. G., Kristensen, T. N., and Loeschcke, V. (2003). The evolutionary and ecological role of heat shock proteins. Ecol. Lett., 1025–1037. doi:10.1046/j.1461-0248.2003.00528.x.

Sun, Z., Kumar, D., Cao, G., Zhu, L., Liu, B., Zhu, M., et al. (2017). Effects of transient high temperature treatment on the intestinal flora of the silkworm Bombyx mori. Sci. Rep. 7, 3349. doi:10.1038/s41598-017-03565-4.

Therneau, T. M. (2020). A package for survival analysis in R. R package version 3.2-7, URL: https://CRAN.R-project.org/package=survival>.

Tôrres, N. M., and Diniz-Filho, J. A. F. (2004). Phylogenetic autocorrelation and evolutionary diversity of Carnivora (Mammalia) in Conservation Units of the New World. Genet. Mol. Biol. 27, 511–516. doi:10.1590/S1415-47572004000400008.

Walters, A. W., Hughes, R. C., Call, T. B., Walker, C. J., Wilcox, H., Petersen, S. C., et al. (2020). The microbiota influences the Drosophila melanogaster life history strategy. Mol. Ecol. 29, 639–653. doi:10.1111/mec.15344.

Wernegreen, J. J. (2012). Mutualism meltdown in insects◻: bacteria constrain thermal adaptation. Curr. Opin. Microbiol. 15, 255–262. doi:10.1016/j.mib.2012.02.001.

Zhang, B., Leonard, S. P., Li, Y., and Moran, N. A. (2019). Obligate bacterial endosymbionts limit thermal tolerance of insect host species. Proceeding Natl. Acad. Sci. 116, 24712–24718. doi:10.1073/pnas.1915307116.

Ziegler, M., Seneca, F. O., Yum, L. K., Palumbi, S. R., and Voolstra, C. R. (2017). Bacterial community dynamics are linked to patterns of coral heat tolerance. Nat. Commun. 8, 14213. doi:10.1038/ncomms14213.

